# Development of a pest threshold decision support system for minimising damage to winter wheat from wheat bulb fly, *Delia coarctata*

**DOI:** 10.1101/2021.03.13.435242

**Authors:** Daniel J Leybourne, Kate E Storer, Pete Berry, Steve Ellis

**Author notes:** These authors contributed equally (Daniel J Leybourne, Kate E Storer).

## Abstract

In this article we describe two predictive models that can be used for the integrated management of wheat bulb fly. Our first model is a pest level prediction model and our second model predicts the number of shoots a winter wheat crop will achieve by the terminal spikelet developmental stage. We revise and update current wheat bulb fly damage thresholds and combine this with our two models to devise a tolerance-based decision support system that can be used to minimise the risk of crop damage by wheat bulb fly.

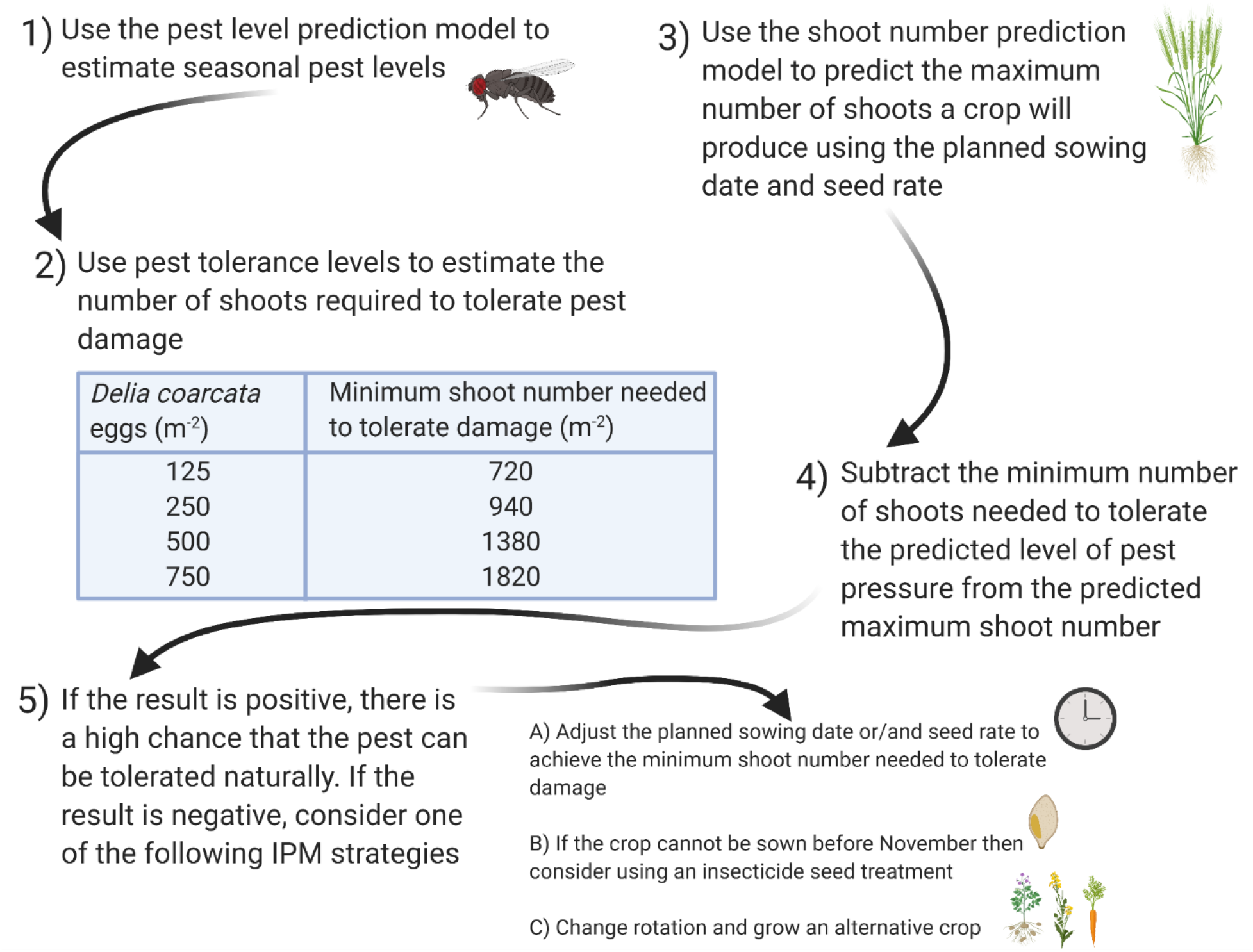

**Summary:** Wheat bulb fly, *Delia coarctata,* is an important pest of winter wheat in the UK, causing significant damage of up to 4 t ha^-1^. Accepted population thresholds for *D. coarctata* are 250 eggs m^-2^ for crops sown up to the end of October and 100 eggs m^-2^ for crops sown from November. Fields with populations of *D. coarctata* that exceed the thresholds are at higher risk of experiencing economically damaging pest infestations. In the UK, recent withdrawal of insecticides means that only a seed treatment is available for chemical control of *D. coarctata,* however this is only effective for late-sown crops (November onwards) and accurate estimations of annual population levels are required to ensure a seed treatment is applied if needed. As a result of the lack of post-drilling control strategies, the management of *D. coarctata* is becoming increasingly reliant on non-chemical methods of control. Control strategies that are effective in managing similar stem-boring pests of wheat include sowing earlier and using higher seed rates to produce crops with more shoots and greater tolerance to shoot damage.

In this study we develop two predictive models that can be used for integrated *D. coarctata* management. The first is an updated pest level prediction model that predicts *D. coarctata* populations from meteorological parameters with a predictive accuracy of 70%, which represents a significant improvement on the previous *D. coarctata* population prediction model. Our second model predicts the maximum number of shoots for a winter wheat crop that would be expected at the terminal spikelet development stage. This shoot number model uses information about the thermal time from plant emergence to terminal spikelet, leaf phyllochron length, plant population, and sowing date to predict the degree of tolerance a crop will have against *D. coarctata.* The shoot number model was calibrated against data collected from five field experiments and tested against data from four experiments. Model testing demonstrated that the shoot number model has a predictive accuracy of 70%. A decision support system using these two models for the sustainable management of *D. coarcata* risk is described.

## Introduction

The wheat bulb fly, *Delia coarctata* (Fallén), is an important herbivorous insect of wheat in the UK. Significant economic damage to winter wheat crops is caused by *D. coarctata* larvae between January and April (Fig. 1) when larvae infest the developing shoots of cereal crops, causing shoot discolouration and stunting (‘deadhearts’). Economic damage can vary between years, reaching 4 t h^-1^ in years of significant infestation (Rogers et al., 2014). *D coarctata* larvae feed until late spring before pupating at the base of the plant, upon emergence adult *D. coarctata* feed on saprophytic fungi present on plant tissue (Jones, 1970) and reproduce before migrating to adjacent fields where oviposition occurs on the bare soil (Bardner et a., 1977).

**Fig. 1:**
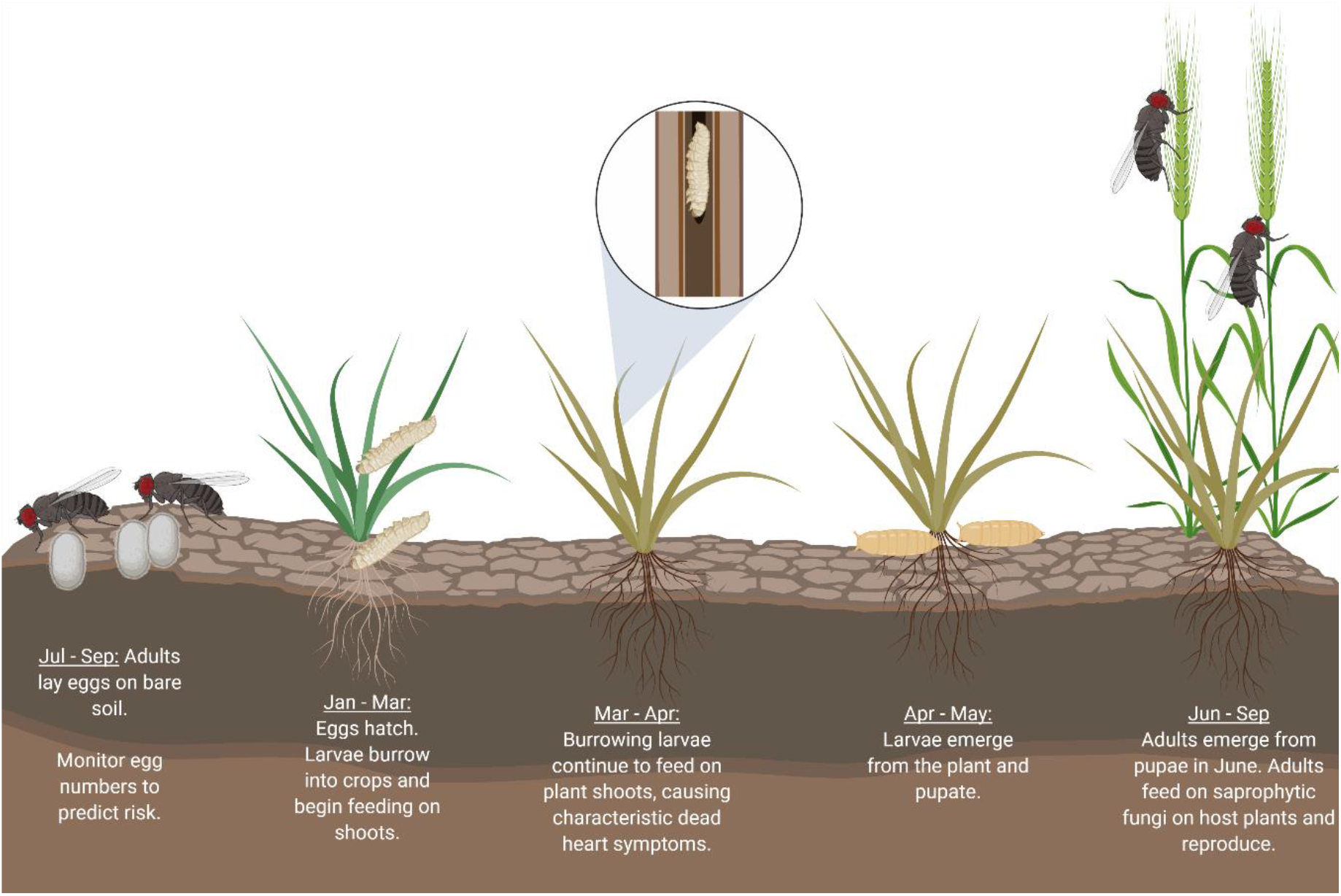
Illustrative example of the annual life cycle of D. coarctata – this image was created with BioRender.com

The level of *D. coarctata* risk fluctuates yearly (Young & Cochrane, 1993). Previous studies have indicated that this correlates with January temperature, July temperature, and August rain days and can be affected further by the previous crop grown in the rotation and the date of harvest (Young & Cochrane, 1993). In the UK, recent withdrawal of insecticides means that only a seed treatment (Signal 300 ES 300 g/l cypermerthrin, UPL Europe Ltd) is available for chemical control of *D. coarctata.* This seed treatment is only effective for late-sown crops (November onwards) as the active ingredient is insufficiently persistent for crops sown at the more conventional drilling dates of September and October. Therefore, alternative non-chemical means of *D. coarctata* control, that are capable of managing *D. coarctata* in crops sown at conventional drilling dates, are becoming increasingly desirable.

One option for the non-chemical control of *D. coarctata* is to adjust crop management practices in order to produce a wheat crop that is able to tolerate *D. coarctata* damage. Crop tolerance, or the economic injury level, can be broadly defined as the amount of pest damage that a crop can withstand before an economic consequence is observed (Stern et al., 1959). The most recent *D. coarctata* pest thresholds of 250 or 100 eggs m^-2^, for crops sown before the end of October and from November, respectively, were devised *c.* 60 years ago by Gough et al (1961) and only take account of pest abundance with no consideration of crop tolerance. Adjusting various agronomic factors, such as sowing date and seed rate, has been used to successfully achieve tolerance against other stem-boring pests of wheat, including the gout fly, *Chlorops pumilionis* (Bjerkander) (Bryson et al., 2005) and wheat saw fly, *Cephus cinctus* Norton (Beres et al., 2011). Modifying agronomic practices to achieve crop tolerance represents a potential method of non-chemical *D. coarcata* control, but to achieve this accurate *D. coarctata* thresholds are required.

For stem-boring insects, crop tolerance can be achieved by growing a crop with a greater number of shoots than those required to achieve an economically viable yield, this allows some shoots to be lost to insect herbivory without incurring a yield penalty. It has been shown that UK wheat crops require a minimum of 400 to 450 fertile shoots (ears) m^-2^ to achieve a typical commercial grain yield (Spink et al., 2000). Wheat crops typically produce more than 1,000 shoots m^-2^ (and up to 1,600 shoots m^-2^) by the start of stem extension in March (Berry et al., 2003). From March to May there is a decline in shoot numbers as the weakest shoots die, leaving a final shoot number of 400 to 700 fertile shoots m^-2^. In most cases, the majority of the shoots are produced during the autumn months, and are therefore present when *D. coarcata* larvae infest plant tissue (January – April; Fig. 1). The main crop management methods that can be used to increase maximum shoot number in spring are to sow early in autumn, allowing more time for extra tillers to develop, and/or to sow a high rate of seeds so that more plants establish, resulting in more shoots m^-2^ (Spink et al., 2000). Whilst it is well understood that earlier sowing and higher seed rates usually result in more shoots m^-2^ (Spink et al., 2000; Darwinkel et al., 1977), reliable methods for quantifying how many more shoots, or how tolerant a crop will be to to *D. coarcata,* do not exist. It should also be recognised that manipulating crop management to produce more shoots often has a practical or economic cost. For example, early sown crops are more prone to lodging and it is not possible to sow early following late harvested root crops. In addition, adverse weather may prevent early sowing and higher seed rates will increase seed costs.

Annual *D. coarctata* pest levels are predicted through soil sampling surveys carried out in September and October, with *D. coarctata* eggs extracted and counted from the sampled soil. This process is time-intensive, the assessment of one sample can take up to three hours (Ramsden, et al., 2017), and requires a suite of bulky equipment (Ramsden et al., 2017; Salt & Hollick, 1944) which are often only available in specialist analytical laboratories. An efficient alternative means of predicting *D. coarctata* pest levels is through predictive modelling (Young & Cochrane, 1993). The Young & Cochrane *D. coarctata* population level prediction model is based on egg counts for East Anglia, UK between 1952 and 1991 (Young & Cochrane, 1993). The Young & Cochrane model predicts *D. coarctata* egg numbers using a range of meteorological parameters, including the departure from the long-term average for rainfall during October of the preceding year, January air temperature, January soil temperature, and July air temperature. Meteorological parameters used in the Young & Cochrane (1993) model were selected based on the hypothesis that they influence either the reproductive development or oviposition of the emerging *D. coarctata* generation. The model developed by Young & Cochrane has a predictive accuracy of 59%, although this is a satisfactory level of prediction for a pest prediction model (Yonow et al., 2004) there is scope to refine the model to further increase its accuracy and reliability. Accurate and reliable prediction of annual *D. coarctata* pest levels before crops are sown is essential if agronomic practices are going to be adjusted to achieve successful *D. coarctata* control. Accurate estimates of plant populations/shoot numbers are also required if practices such as early sowing or sowing at a higher seed rate are going to be used to improve crop tolerance to *D. coarctata,* as has been the case for similar wheat stem-borers (Beres et al., 2011; Bryson et al., 2005).

Here, we develop two predictive models that can be used for integrated *D. coarctata* control. The first is an updated pest level prediction model that estimates *D. coarctata* populations from meteorological parameters with a greater predictive accuracy than the previous model developed by Young and Cochrane (1993). The second is a model that predicts the maximum number of shoots for a winter wheat crop just prior to the start of stem extension based on target plant population and sowing date; this will be particularly valuable when trying to optimise the target plant population and sowing date to produce crops which are able to tolerate *D. coarctata* infestation. Finally, using data extracted from the literature we produce a revised calculation of *D. coarctata* threshold levels and use this to develop a sustainable decision support system for minimising the risk of economic crop damage by *D. coarctata.*.

## Methods

### Modelling to predict D. coarctata egg numbers

#### *Sources of* D. coarctata *egg numbers and meteorological data*

*Delia coarctata* egg number data were extracted from two sources. Historic data from East Anglia (1952 – 1991) were extracted from Young & Cochrane (1993) and combined with the results from the AHDB Autumn survey of *D. coarctata* incidence from northern England (2005 – 2019) and East Anglia (2008 – 2019). Up to 30 fields were sampled in September or October of each survey year in areas prone to *D. coarctata* infestation, with *c.* 15 in eastern England and 15 in northern England. Samples were taken in September or October once egg laying was complete (usually early September; Fig. 1). For each field sampled, either 32 cores (each of 7.2 cm diameter) or 20 cores (each of 10 cm diameter) were taken to cultivation depth. Fields were sampled in a standard ‘W’ sampling pattern across the direction of cultivation. *D. coarctata* eggs were extracted following soil washing (Salt & Hollick, 1944) and flotation in saturated magnesium sulphate. Egg numbers were expressed as number of eggs m^-2^. Meteorological data were extracted from UK meteorological office data for each region (www.metoffice.gov.uk/research/climate/maps-and-data/uk-and-regional-series). Meteorological data extracted included minimum, mean, and maximum temperature, rain days, rainfall amount (mm), sunshine days, and air frost days. From the extracted data, the deviation from the long-term average was calculated. Long-term averages of 30-year periods were used, the only deviation for this was for air frost for which reporting did not commence until 1960. The UK meteorological office categorises seasons into: Winter (preceding December – February), spring (March – May), summer (June – August), autumn (September – November). For each year of egg collection, the meteorological data included in the model were the autumn of the preceding calendar year, winter (starting in December of the preceding year), current spring, and current summer. For each season the meteorological parameters included were: minimum temperature, mean temperature, maximum temperature, the number of sun days, the number of rain days, and the amount of rainfall. The number of air frost days was included for the winter, spring, and preceding autumn seasons only.

#### Modelling approach – developing the pest level prediction models

Data modelling for the pest level prediction model was carried out in R v.3.6.1 with additional package ggplot 2 v.3.2.1 (Wickham, 2016) used for data visualisation. Linear regressions were used to build all models and backwards stepwise model selection was employed to arrive at the final predictive models (Marill & Green, 1963). At each simplification stage analysis of variance was carried out to ensure model simplification was justified and did not significantly affect model structure. Model residuals were observed at each stage.

#### Seasonal and monthly models for 1952 – 2019

An initial seasonal model was developed using the following meteorological factors on a seasonal basis: minimum air temperature, mean air temperature, maximum air temperature, rain days, rainfall, and sun days. The model was refined through backwards stepwise model selection until the final seasonal model was produced. Based on this final seasonal model an initial monthly model was developed that included monthly inputs for all the parameters included in the final seasonal model (e.g. the final seasonal model included summer minimum temperature, therefore the initial monthly model included minimum temperature for June, July, and August). The monthly model was simplified through backward stepwise model selection until the final monthly model was produced.

#### Seasonal and monthly models for 1971 – 2019 (air frost models)

In order to allow air frost to be included in the model a subdataset was developed comprising all observations from 1971 – 2019. The 1971 – 2019 period used in this subdataset was to allow the long-term average to be calculated for a minimum period of ten-years. Seasonal and monthly models were simplified as described above and the final model was used to predict egg numbers.

The final monthly model was validated following a similar procedure to the one employed by Young & Cochrane (1993): four five-year periods were removed from the dataset from which the model was developed and observations were made of how this affected the model predictions. Data were removed from the first five years (validation 1: years 1971 – 1975 removed), the last five years (validation 2: years 2015 – 2019), the five years with the highest recorded *D. coarctata* egg numbers (validation 3: years 1978, 1984, 1985, 1986, 2010), and the five years with the lowest recorded *D. coarctata* egg numbers (validation 4: years, 2005, 2006, 2007, 2014, 2017). A further four validations (validations 5-8) were carried out by randomly removing five one-year periods from the dataset from which the model was developed.

Soil samples were taken from 30 sites in England (divided into 15 northern and 15 eastern sites) in September and October 2020 and the number of *D. coarctata* eggs per sample were determined; these samples were used to independently test the prediction model.

### Shoot number model

#### Model of potential shoot number

Data modelling for the shoot number prediction model was carried out in Microsoft Excel and involved two separate processes. First, the potential shoot number of a single wheat plant growing in isolation (i.e. without any competition from neighbouring plants and unlimited resources) was determined. Secondly, this calculation was calibrated using field data to account for plant competition and environmental factors that might limit shoot production.

#### Building a thermal time-based model for shoot production

Published principles of wheat shoot development were used to develop a thermal time-based model for shoot production (Klepper et al., 1984). The thermal duration between sowing and plant emergence was taken as 150°Cd (Sylvester-Bradley et al., 1998). The end of tillering generally coincides with the start of stem extension and the formation of the terminal spikelet within the developing ear. The thermal time between sowing date and terminal spikelet production for early and late sown winter wheat crops was reported in Kirby et al. (1999). Data from Kirby et al. (1999) was used to estimate the effect of sowing date (1^st^ September to 8^th^ November) on the thermal duration between sowing and terminal spikelet, assuming that it decreased linearly with time of sowing. This meant that the thermal time from sowing to terminal spikelet decreased by 9°C for each day that sowing was delayed after 1^st^ September from a value on 1^st^ September of 1582°Cd. The thermal time from sowing to plant emergence was then subtracted from this value, leaving the total thermal time available for leaf and shoot production.

The thermal time between emergence of successive leaves (phyllochron) decreases the later crops are sown (Equation 1) where d = the number of days after 1st September when the crop was sown (Kirby et al., 1985).

Phyllochron lengh = −0.5383 × *d* + 140.3

**Equation 1**

The phyllochron length, along with the thermal time between plant emergence and terminal spikelet, was then used to estimate the number of leaves and shoots that could be produced over time for an individual plant. The following assumptions were made based on Klepper et al., (1984):

- Three phyllochrons after plant emergence: The first primary shoot emerges from the axil of the 1^st^ leaf on the main shoot
- Four phyllochrons after plant emergence: The second primary shoot emerges from the axil of the second leaf of the main shoot
- Five phyllochrons after plant emergence: The third primary shoot emerges from the axil of the third leaf of the main shoot. The first secondary shoot emerges from the axil of the first leaf of the first primary shoot
- Six phyllochrons after plant emergence: The fourth primary shoot emerges from the axil of the fourth leaf of the main shoot. The second secondary shoot emerges from the axil of the second leaf of the first primary shoot. The third secondary shoot emerges from the axil of the first leaf of the second primary shoot.

#### Calibrating and testing the shoot number model

Data from five winter wheat trials (Table S1) were used to calibrate the shoot number model and data from three winter wheat trials (Table S2), alongside data from Spink et al., 2000, were used to test the shoot number model. Each of the field trials (Table S1; S2) used the same winter what variety (Evolution) and included a range of seed rates (40, 80, 160, 320, 480, and 640 seeds m^-2^). All trials were sown at either ‘standard’ or ‘late’ timings for the region, and were treated with an insecticide to control for *D. coarctata* (Tables S1; S2). The experiment at Huggate 2016 was not treated with insecticide and assessments showed that less than 1% of shoots were infested with *D. coarctata* at the start of stem extension. Each experiment was arranged in a fully randomised block design with either three or four replicates of each treatment (Tables S1; S2). Experimental plots were 2m x 12m and were drilled using an Ojyard experimental plot drill. Plant number and shoot number were measured at the start of stem extension (BBCH Growth Stage 31 (GS31)) in each experimental plot by counting all plants and shoots within a 0.7m x 0.7m quadrat. Field trial data (Table S1) were analysed in Genstat (v-14) using a one-way ANOVA. Standard error of the difference (SED) values are reported alongside p-values where relevant. Data used to calibrate the model were not used for model testing. The model was tested against three field trials (Table S2) and data extracted from a previous seed rate and sowing date winter wheat experiment (Spink et al., 2000). Multiple linear regression analysis, with Experiment included as ‘Group’, was used to compare the predicted shoot numbers against the model predictions using Genstat (v-14).

## Results

### *The* D. coarctata *pest level prediction model*

In order to improve on a previous *D. coarctata* population prediction model (the Young & Cochrane 1993 model), open-access meteorological data (published by the UK Meteorological Office) were extracted on a seasonal and monthly basis and incorporated into two linear models. In order to identify which seasons are most important in determining *D. coarctata* egg numbers an initial model was developed on a seasonal basis. This model (1952 – 2019 seasonal model; Fig. S1A) indicated that the number of preceding autumn rain days, the minimum winter temperature, spring mean temperature, spring maximum temperature, spring rainfall, and summer minimum temperature are the most important meteorological parameters affecting *D. coarctata* egg numbers on a seasonal basis (adjusted R^2^ = 0.49; F_6,60_ = 11.54; p = <0.001). The seasonal meteorological parameters listed above were used as inputs for the predictive model. Model predictions versus observations for the seasonal 1952 – 2019 model, including model predictions for 1992 – 2004, are displayed in Fig. S1A.

In order to identify the months which influence *D. coarctata* egg numbers the meteorological parameters included in the seasonal 1952 – 2019 model were assessed on a monthly basis for the relevant seasons: preceding September – preceding November for preceding autumn season, preceding December – February for winter season, and June – August for summer season. Linear regression modelling indicated that the most important monthly meteorological parameters for predicting *D. coarctata* egg numbers were preceding October rain days, preceding December minimum temperature, January minimum temperature, March mean temperature, May mean temperature, March maximum temperature, May maximum temperature, May rainfall, June minimum temperature, July minimum temperature, and August minimum temperature (F_11,55_ = 6.66; p = <0.001; adjusted R^2^ = 0.49). The monthly meteorological parameters listed above were used as inputs for the predictive model. The predictions of this model, compared with the observed values, are shown in Fig. 2A.

**Fig. 2:**
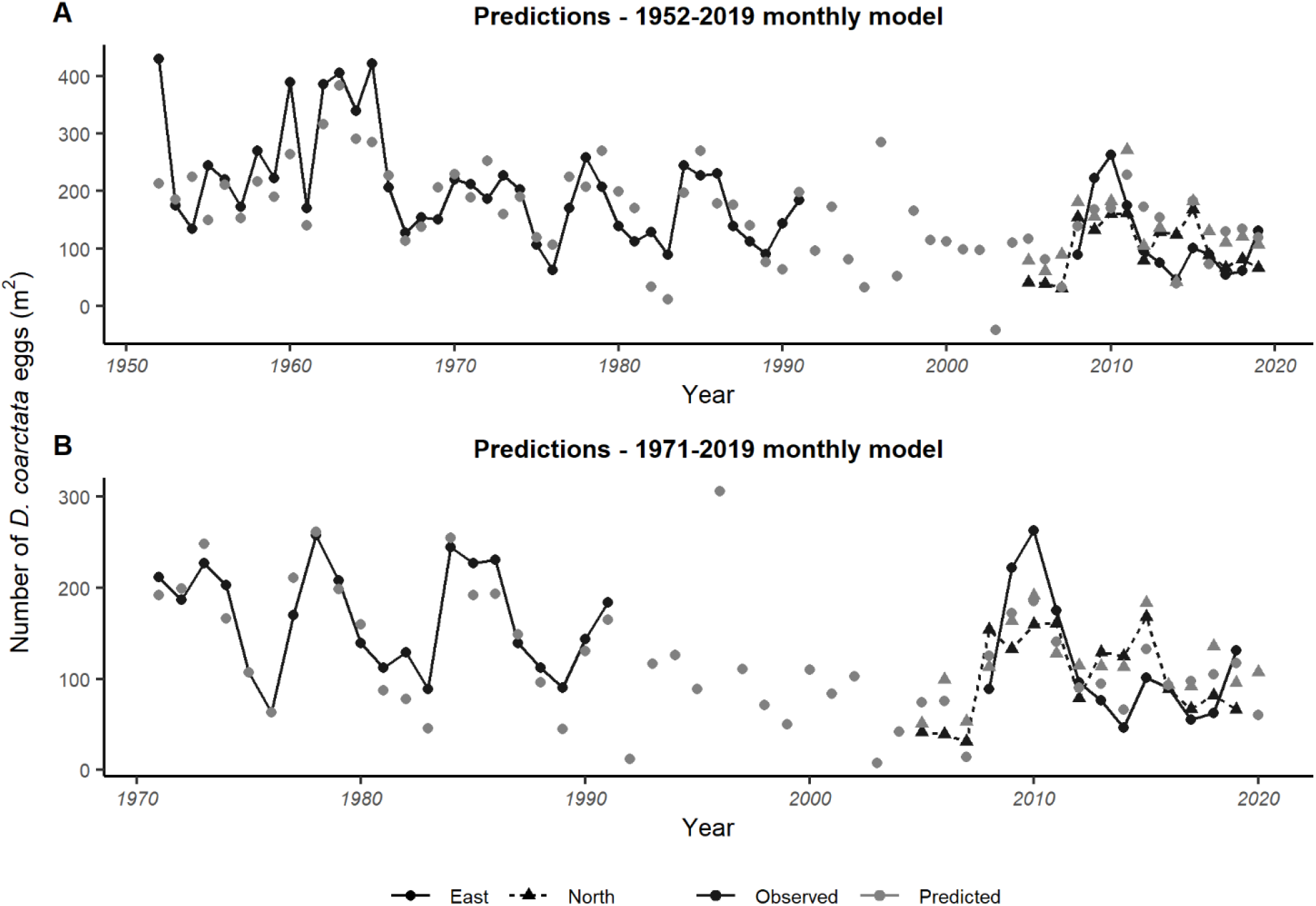
Predicted and observed D. coarctata egg numbers. A) 1952 – 2019 monthly model, B) 1971 – 2019 monthly model. Predictions (grey) are plotted alongside the mean observed value (black) and divided into the two regions, north (triangle; dashed line) and east (circle; solid line). For clarity, trendlines are included for the observed values only.

### Introducing air frost into the pest prediction model increases predictive power

One key factor that might influence *D. coarctata* egg numbers is air frost. The UK meteorological office began long-term monitoring of air frost in 1960. A seasonal model was produced using data from the 1971 – 2019 subdataset. This comprised of all meteorological parameters described above as well as the departure from long-term average for preceding autumn, winter, and spring, air frost days. Following model simplification, this air frost seasonal model had a higher predictive power compared with the previously developed models (adjusted R^2^ = 0.59, F_9,38_ = 842; p = <0.001) and the final meteorological parameters included departure from long-term average for: preceding autumn rain days, preceding autumn sun days, winter mean temperature, winter air frost days, spring maximum temperature, spring rainfall, summer minimum temperature, summer mean temperature, and summer maximum temperature. These seasonal meteorological parameters were used as inputs for the predictive model. The predictions of this model, compared with the observed values, are shown in Fig. S1B.

A refined 1971 – 2019 monthly model was created using monthly average data for the seasons shown to be important in the seasonal model, using the same approach described above for the 1952-2019 model. The meteorological inputs for this refined monthly model were the departure from long-term average for: preceding September sun days, preceding October rain days, January mean temperature, January frost, April maximum temperature, May maximum temperature, April rainfall, and July minimum temperature. This model had the highest predictive power (adjusted R^2^ = 0.70, F_8,39_ = 14.88; p = <0.001). The monthly meteorological parameters listed above were used as inputs for the predictive model. The predictions of this model, compared with the observed values, are shown in Fig. 2B. On average, the predicted values only deviated from the observed values by −9% (median = −2%; range = −155% to +50%).

### Validating the final pest prediction model

The 1971 – 2019 monthly model was validated by removing a series of years from the model, rerunning the model, and observing the effect the removal of these years had on the ability of the model to predict *D. coarctata* egg numbers for all years (1971 – 2019), similar to the validation process deployed by Young & Cochrane (1993). Eight validation models were developed in total (Fig. S2). Validations had no significant detrimental effect on the predictive power of the models; average deviation from the predictive values of the full model were: −20.20% (validation 1), +0.05% (validation 2), +0.58% (validation 3), −17.10% (validation 4), +2.72% (validation 5), −1.28% (validation 6), +2.70% (validation 7), −1.94% (validation 8). The relationship between the observed and predicted values of the final 1971 – 2019 monthly model is shown in Fig. S3.

### *Testing the* D. coarctata *prediction model*

The model was used to predict mean *D. coarctata* egg numbers for each region in 2020. The model predictions were then compared with the mean egg counts per region obtained by soil sampling. The model predicted a mean *D. coarctata* egg number of 60 eggs m^-2^ for eastern England and 107 eggs m^-2^ northern England. Soil sampling (15 sites per region) indicated that the observed regional risk was 111 eggs m^-2^ for northern England, and 173 eggs m^-2^ for eastern England. Higher observed values for eastern England than were estimated were mainly driven by three sites with very high counts of 1000, 850, and 404 eggs m^-2^.

### Predicting shoot production number for individual plants growing in isolation

The shoot production model estimated the shoot production of a single plant grown in isolation at a range of sowing dates between 1st September and 30th December (Fig. 3). The model predicted that a single plant sown on 1st September has the potential to produce 38 shoots by terminal spikelet (which approximates to the start of stem extension), whereas at the other extreme a single plant sown in mid-November would only produce ten shoots.

**Fig. 3:**
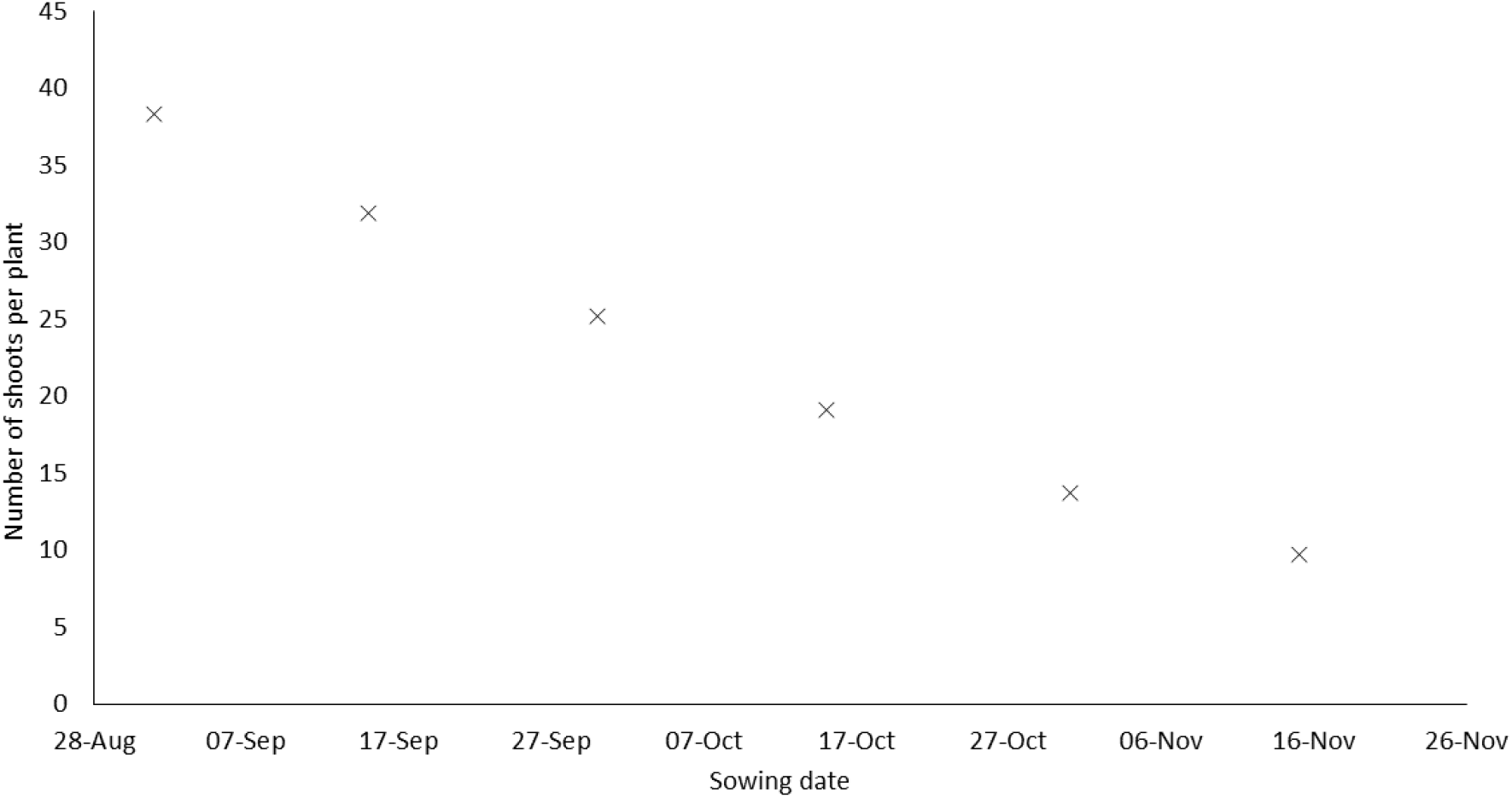
Predicted number of shoots per plant produced by terminal spikelet (which approximates to the start of stem extension), for a given sowing date when grown in the absence of competition and environmental stress factors.

### Calibrating the shoot production number model

It was recognised that achieving a shoot population of 38 shoots per plant (Fig. 3) was unrealistic under field conditions, primarily due to competition between shoots for limited resources but also due to stress factors (e.g. soil capping, pests, disease). Shoot number data from specific treatments in a series of field trials (Table S1) were used to calibrate the model to ensure that it provided a realistic estimation for the number of shoots that could be expected in field conditions. The ratio of observed to predicted shoots per plant was negatively related to the observed plants m^-2^ (Fig. 4). This was because the shoot number model predicted the highest potential number of shoots m^-2^ for high plant populations, but these populations also had the greatest competition between shoots resulting in the lowest ratio of observed to predicted shoots per plant. The relationship between the observed plants m^-2^ and ratio of observed to predicted shoots per plant described in Fig. 4 was used to calibrate the potential shoot number model for field conditions.

**Fig. 4:**
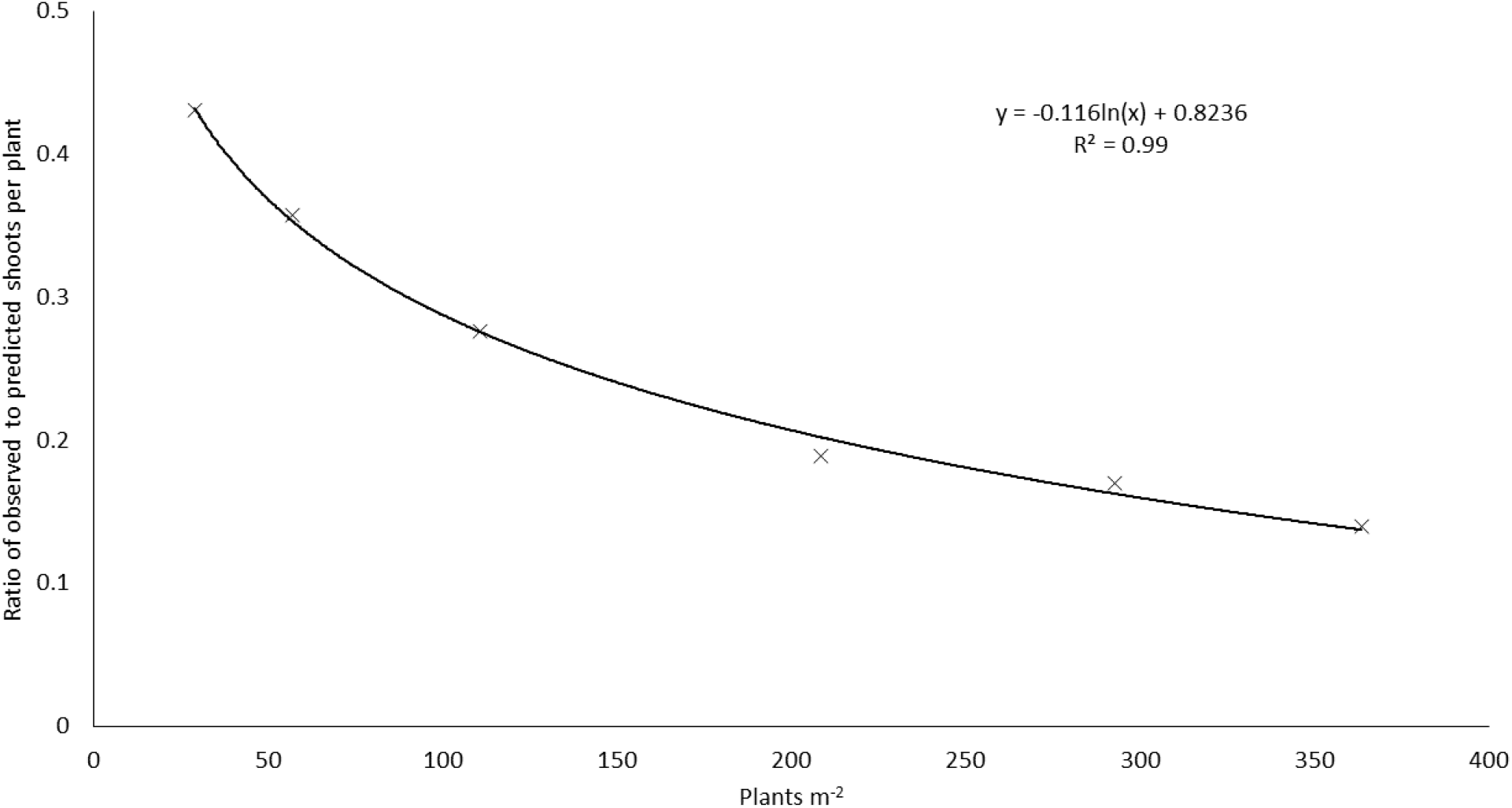
The ratio of observed to predicted shoots per plant at the beginning of stem extension plotted against measured plants m^-2^ for crops sown in the first week of October.

The calibrated shoot number model predicts a decline in maximum shoot number due to reduced plant population and delayed sowing date, as is generally observed in practice (Fig. 5). A crop sown at the end of September with a plant population of 200 plants m^-2^ is predicted to produce a maximum shoot number of approximately 1071 shoots m^-2^. The model can be used to estimate the minimum plants m^-2^ required to achieve 500 shoots m^-2^ (the minimum number of shoots required to achieve a typical commercial UK wheat yield). To achieve 500 shoots m^-2^ by terminal spikelet the model estimates that a minimum of 56 plants m^-2^ for late September sowing, 91 plants m^-2^ for mid-October sowing, 163 plants m^-2^ for late October sowing, and 418 plants m^-2^ for mid-November sowing are required. These plant populations are similar, or slightly greater, than estimates of the economic optimum plant density reported by Spink et al. (2000), which provides confidence that the shoot number model is giving plausible predictions. Although the number of shoots can further increase between terminal spikelet stage and harvest, the damage caused by *D. coarctata* occurs prior to terminal spikelet (Fig. 1). Therefore, shoot production up to terminal spikelet was considered most appropriate for developing a *D. coarctata* tolerance scheme. The shoot production model has been used to quantify the increase in maximum shoot number by terminal spikelet as a result of sowing earlier and establishing a higher plant population (Fig 6). Sowing earlier generally results in a larger increase in shoots m^-2^ compared with increases in plant population. This information can be used to help estimate changes in sowing date and seed rate to minimise the risk of yield loss to *D. coarcata.*

**Fig. 5:**
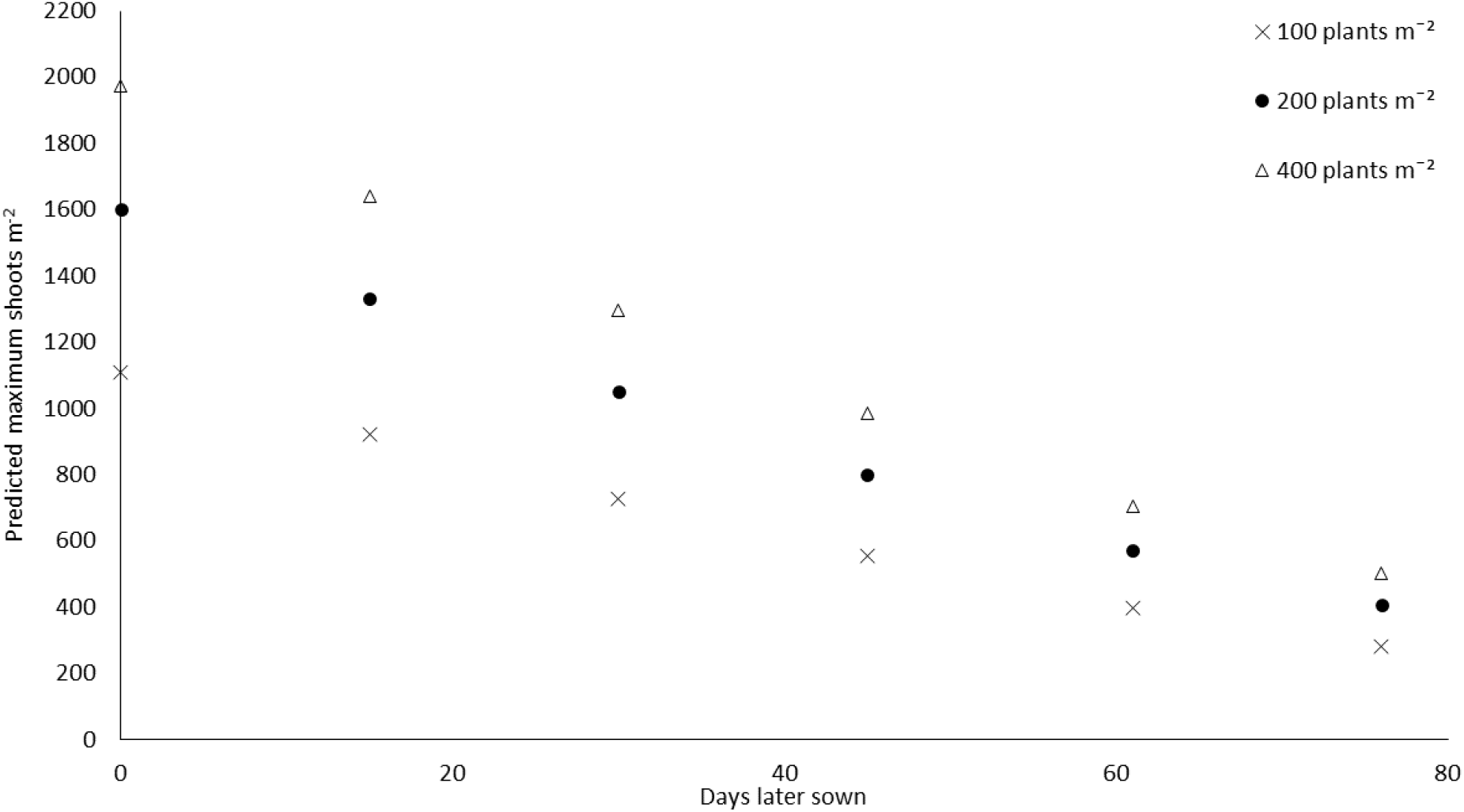
Number of shoots m^-2^ predicted by the shoot number model after calibration for field conditions for crops sown on different sowing dates after 1^st^ September with plant populations of 100, 200, and 400 plants m^-2^.

**Fig. 6:**
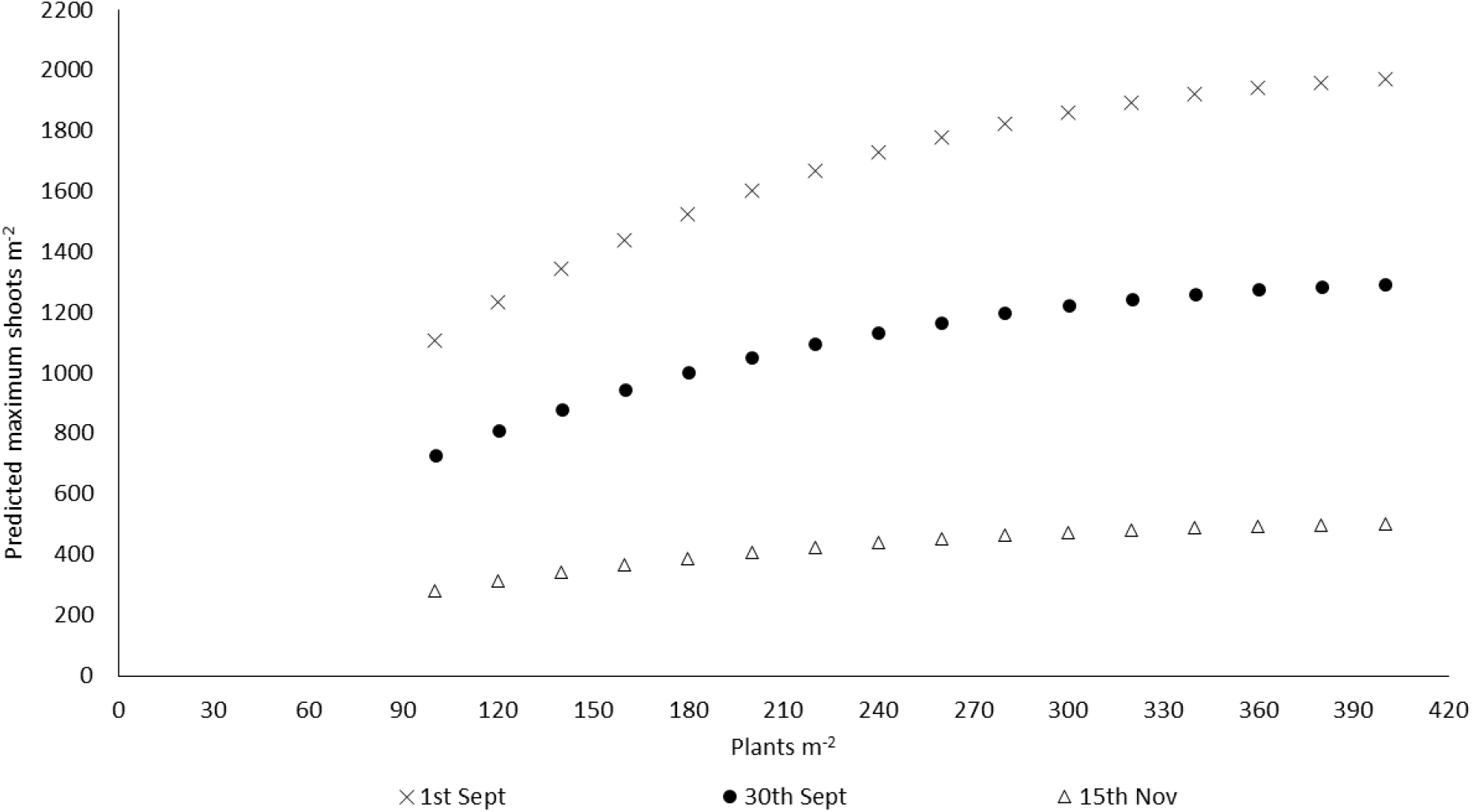
Effect of changes in sowing date and plant population on the predicted number of shoots m^-2^.

### Testing the shoot number prediction model

Data from three winter wheat field experiments (Table S2) were combined with data from other field experiments (Spink et al., 2000) to test the predictive power of the model. (Fig. 7). The number of shoots m^-2^ at GS31 were measured in each of the field trials (Table S2), which approximates to the timing of terminal spikelet production. Multiple linear regression analysis showed that a single best fit line had an R^2^ value of 0.70 (p = <0.001) (Fig. 7). Fitting separate best fit lines to each experimental site with the same slope but different y axis intercepts gave an R^2^ value of 0.83 (p = <0.001). Across all the experimental sites the single best fit line was slightly greater than the 1:1 relationship.

**Fig. 7:**
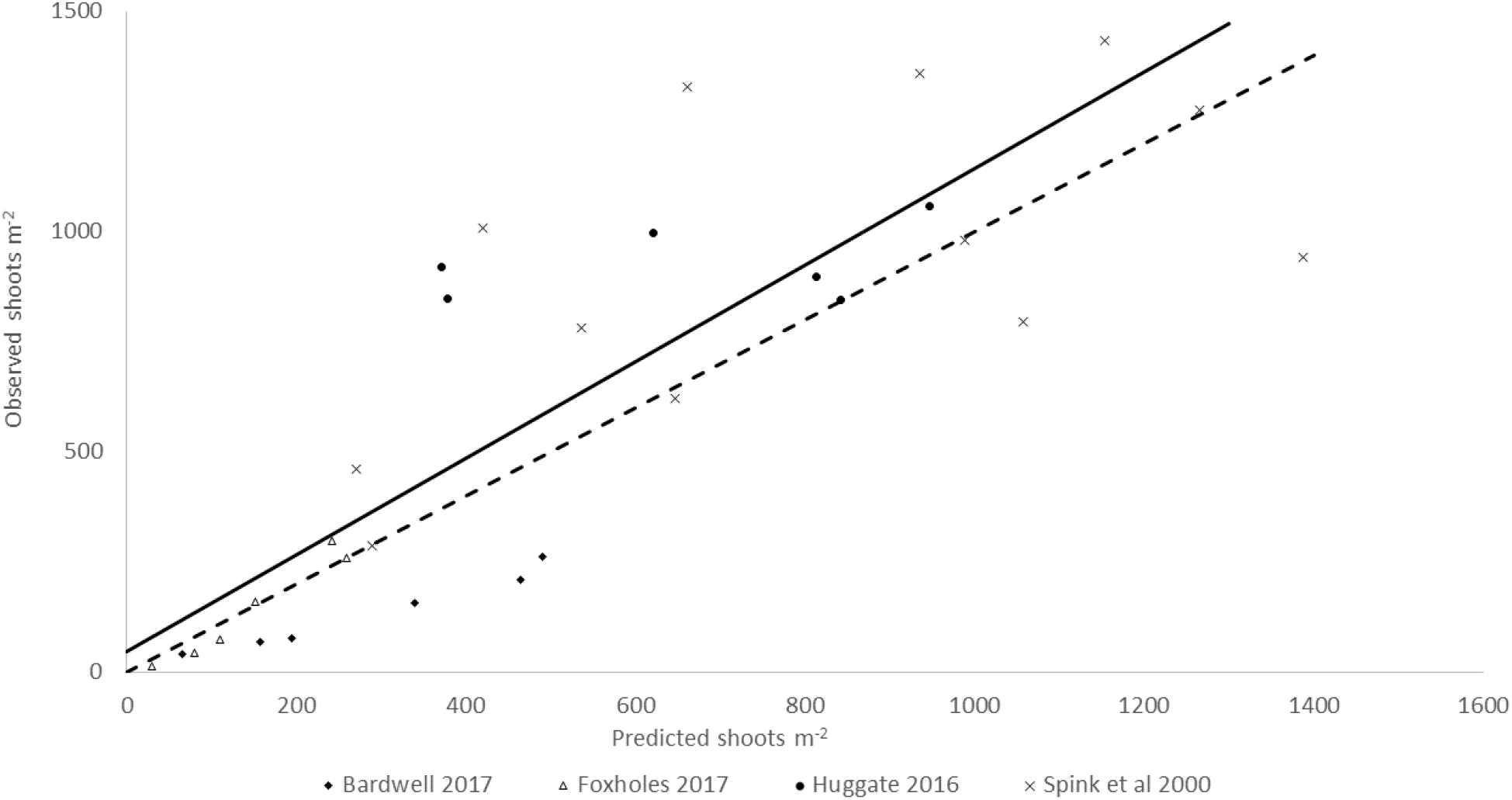
Model predicted shoots m-2 plotted against the observed shoots m^-2^ for data measured in the experiments listed in Table S2 and Spink et al., 2000. Solid black line represents the linear regression fitted to all data (R^2^ = 0.70). The dashed line represents a 1:1 relationship.

### *Revising the economic injury level of wheat to* D. coarctata

The following factors determine how much damage a wheat crop can sustain from a stem-boring insect before the damage becomes economically damaging, and can be used to provide a more comprehensive estimation of economic thresholds for *D. coarctata:*

1. The number of shoots a larva can destroy
2. The minimum number of fertile shoots a crop requires to achieve a yield potential
3. The maximum number of shoots a crop is expected to produce in winter

These factors can be used to revise the *D. coarctata* threshold using Equation 2.

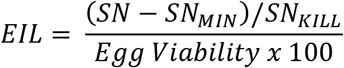

**Equation 2:** Economic Injury Level (EIL) equation used to estimate wheat tolerance against *D. coarctata.* SN = the number of shoots per m^-2^ in winter, SN_MIN_ = the minimum number of fertile shoots per m^-2^ required to achieve a yield potential, SN_KILL_ = the number of shoots killed by an individual larva, and Egg Viability the proportion of eggs that develop into larva.

Using values from the literature the parameters required for the equation can be estimated: Egg viability is estimated at 56% (Ryan, 1973A; Raw, 1967; Gough, 1947), SN_KILL_ is estimated at four shoots destroyed per larva (Young & Ellis, 1996; Ryan, 1975; Ryan, 1973B), SN is estimated at 1000 shoots m^-2^ (Sylvester-Bradley et al. 1998), and SN_MIN_ is estimated at 400-450 shoots m^-2^ (Spink et al., 2000). Using these assumed values, but increasing the SN_MIN_ to 500 shoots m^-2^ to allow a modest degree of insurance against achieving too few shoots, an updated threshold for *D. coarctata* in winter wheat is presented in Table 1 and can be used as the foundation to develop targeted agronomic approaches for minimising the risk of economic damage by *D. coarctata* control using crop tolerance.

**Table 1:** Minimum shoot number at GS31 needed to tolerate different levels of *D. coarctata* damage

| Egg count (eggs m<sup>-2</sup>) | Shoot number m<sup>-2</sup> required to tolerate *D. coarctata* damage |
| --- | --- |
| 125 | 720 |
| 250 | 940 |
| 500 | 1380 |
| 750 | 1820 |

Calculations assume 500 shoots m<sup>-2</sup> is the minimum required to achieve a typical UK commercial wheat yield.

## Discussion

### *Predicting* D. coarctata *risk to inform control measures*

*Delia coarctata* larva infest cereal shoots between January and April, where they can cause devastating crop damage resulting in yield losses of up to 4 t ha^-1^ (Rogers et al., 2014). Depending on the sowing date, current thresholds indicate that a pest pressure of 250 eggs m^-2^ can cause significant crop damage for crops drilled before October, with a pressure of 100 eggs m^-2^ causing significant crop damage for crops drilled from November (Gough, 1961). Chemical-based options for *D. coarctata* control in the UK are limited to a pre-drilling seed treatment. This is only effective for late sown crops (November onwards) and is insufficiently persistent for earlier sowings. Therefore, decisions on whether to apply a seed treatment need to be made before sowing and should take into account the risk of *D. coarctata* infestation. Currently, the most accurate means of determining *D. coarctata* risk requires soil extraction and manual egg counting after the previous crop has been harvested, this is both time- and cost-intensive (Ramsden et al., 2017). Predicting pest levels and risk is an important Integrated Pest Management (IPM) tool and represents a cost-effective means of estimating annual risk for a wide range of important insect pests (Herms, 2004). The pest level prediction model developed in this study provides an alternative to soil sampling and has the significant advantages that it is less time consuming, less arduous, and provides an earlier estimate of pest levels, which is crucial when deciding whether to treat seed. Similar prediction models have been developed for a range of other agriculturally important insect pests, including the tea leaf roller, *Caloptilia theivora* Walsingham (Satake et al., 2005), leafhoppers, whiteflies, and thrips (Arya et al., 2015). Pest level and risk prediction models can predict when a phenological event has occurred that increases the in-season crop risk (such as pest egg hatch, development, or emergence of an additional generation) to help time the application of pest management strategies (Satake et al., 2005; Herms, 2004; Milonas et al., 2001). Alternatively, models can predict annual risk by estimating pest population densities (Arya et al., 2015). As *D. coarctata* can currently only be controlled chemically using a seed treatment, it is important that any pest level prediction model developed can accurately predict pest populations before sowing, in time for sowing date and seed rates to be adjusted.

The *D. coarctata* pest level prediction model developed here is based on the Young & Cochrane model (Young & Cochrane, 1993) but incorporates a wider range of meteorological parameters than the original. The meteorological parameters included in the Young & Cochrane model were selected to include the factors hypothesised to have the greatest influence on *D, coarctata* biology and phenology (Young & Cochrane, 1993). The Young & Cochrane model had a predictive accuracy of 59%, limiting its uptake as a decision support tool for wheat growers. When developing our models, we included additional meteorological factors available from open-access data sources. The benefits of building the models using open-access meteorological data are that the model inputs are standardised across regions and freely available. The meteorological parameters included in the models were departure from the long-term average for: minimum, mean, and maximum temperature, rainfall, the number of rain days (days with rainfall > 0.2 ml), the number of sun days, and days of air frost. Our final model (1971 – 2019 monthly model) had a predictive accuracy of 70%, an 11% increase in accuracy when compared with the Young & Cochrane model. This is a significant improvement of the potential for using predictive modelling to estimate *D. coarctata* risk. Furthermore, as the model can be run prior to sowing in August/September, it is a possible alternative to soil sampling, and it can be used to target seasons and regions where soil sampling should be focussed.

During model testing the model performed well and predicted regional *D. coarctata* risk accurately for the northern region; however, the level of risk predicted for the eastern region was lower than observed. Further model development and refinement (through the potential inclusion of model moderators such as soil type and previous crop) would enable a more robust and dynamic model to be developed. Soil type is likely an important factor to consider in future model development, as the three 2020 test sites with the highest *D. coarctata* counts were associated with clay soils, indicating that soil type might influence *D. coarctata* oviposition preference. Furthermore, the previous crop in the rotation has been reported to affect *D. coarctata* oviposition (Young & Cochrane, 1993; Gough, 1946). Therefore, including these two factors as components in subsequent model improvements represents the logical next step in future model development. The predictive accuracy of our model is similar to the accuracy of other pest prediction models that use meteorological data to predict seasonal risk. Including models that predict the population dynamics of the Queensland fruit fly, *Bactrocera tryoni* Froggatt (R^2^ = 0.28 and 0.32; Yonow et al., 2004), annual populations of the whitetail, *Paronychiurus kimi* (Lee) (R^2^ = 0.79; Choi & Ryoo, 2003), and the abundance of two planthopper species, *Helicoverpa spp.,* (R^2^ = 0.84 – 0.96; Zalucki & Furlong, 2005). Therefore, our model represents a potential useful component of a sustainable IPM strategy.

### *Improving tolerance to* D. coarctata *through shoot number prediction*

Non-chemical methods for *D. coarctata* control are becoming increasingly desirable, from both an agronomic and an environmental perspective. Insecticidal sprays are no longer approved for *D. coarctata* control in the UK and a growing body of evidence indicates that pesticides can have far- reaching environmental consequences (Leather, 2018), resulting in a need to develop non-chemical pest management methods. Cultural control can be an effective means of limiting pest damage (Glen, 2000) and can involve the adjustment of both pre-drilling and in-season agronomic practices. Non-chemical control of similar stem-boring pests of wheat can be effectively achieved through adjustments to pre-drilling agronomic practices, such as increased seed rates and earlier sowings (Glen, 2000). These practices have the potential to increase crop shoot numbers, and therefore improve crop tolerance to herbivorous insects (Wenda-Pieskik et al., 2017; Beres et al., 2011; Bryson et al., 2005).

Higher seed rates have been exploited as an IPM strategy to confer tolerance in wheat against other stem-boring pests, including the wheat stem sawfly, *Ce. cinctus* (Beres et al., 2011) and the gout fly, *Ch. pumilionis* (Bryson et al., 2005). Earlier drilling has also been reported to confer tolerance against cereal leaf beetles, *Oulema spp.* (Wenda-Pieskik et al., 2017). Together, this showcases the potential to achieve cultural control of herbivorous insect pests through the adjustment of pre-drilling agronomic practices, including sowing date and seed rate adjustments. Therefore, accurate means of predicting crop shoot numbers will be an important component of any IPM strategy based on these methods. The shoot number prediction model we developed in this study can be used in conjunction with our revised *D. coarctata* thresholds (Table 1) as a component of an integrative *D. coarctata* risk management system.

Our shoot number model has an overall predictive accuracy of 70% and the predicted values were close to the 1:1 relationship across all the experiments the model was tested against. For some sites the model made more accurate predictions (e.g. Foxholes, 2017), although for others a weaker relationship was observed (e.g. Bardwell, 2017). This contrast in accuracy could be related to specific environmental factors that limited shoot production such as soil capping or water logging (Robertson et al., 2009). Our revised thresholds for *D. coarctata* in winter wheat demonstrate that the current *D. coarctata* pest threshold of 250 eggs m^-2^ (Gough et al., 1961) is too simplistic and for many crops this likely represents either an overestimation of the potential pest damage, an underestimation of the amount of damage that can be tolerated by a winter wheat crop, or a combination of both factors. Sensitivity analysis (achieved by adjusting each parameter used in Equation 2 from its likely minimum value to its maximum value; Fig. 8) demonstrates that thresholds for *D. coarctata* could be smaller or greater than 250 eggs m^-2^, and that the thresholds are particularly sensitive to the number of shoots a crop will produce. Combining the shoot prediction model with the revised *D. coarctata* thresholds will facilitate the production of a winter wheat crop that is capable of tolerating the predicted level of *D. coarctata* damage through adjustments to seed rate, sowing date, or both. This approach is similar to those devised by Bryson et al., (2005), Beres et al., (2011), and Wenda-Pieskik et al., (2017). Although, increasing shoot number would need to be considered alongside higher crop production costs and the increased risk of crop lodging (Berry & Spink, 2012).

**Fig. 8:**
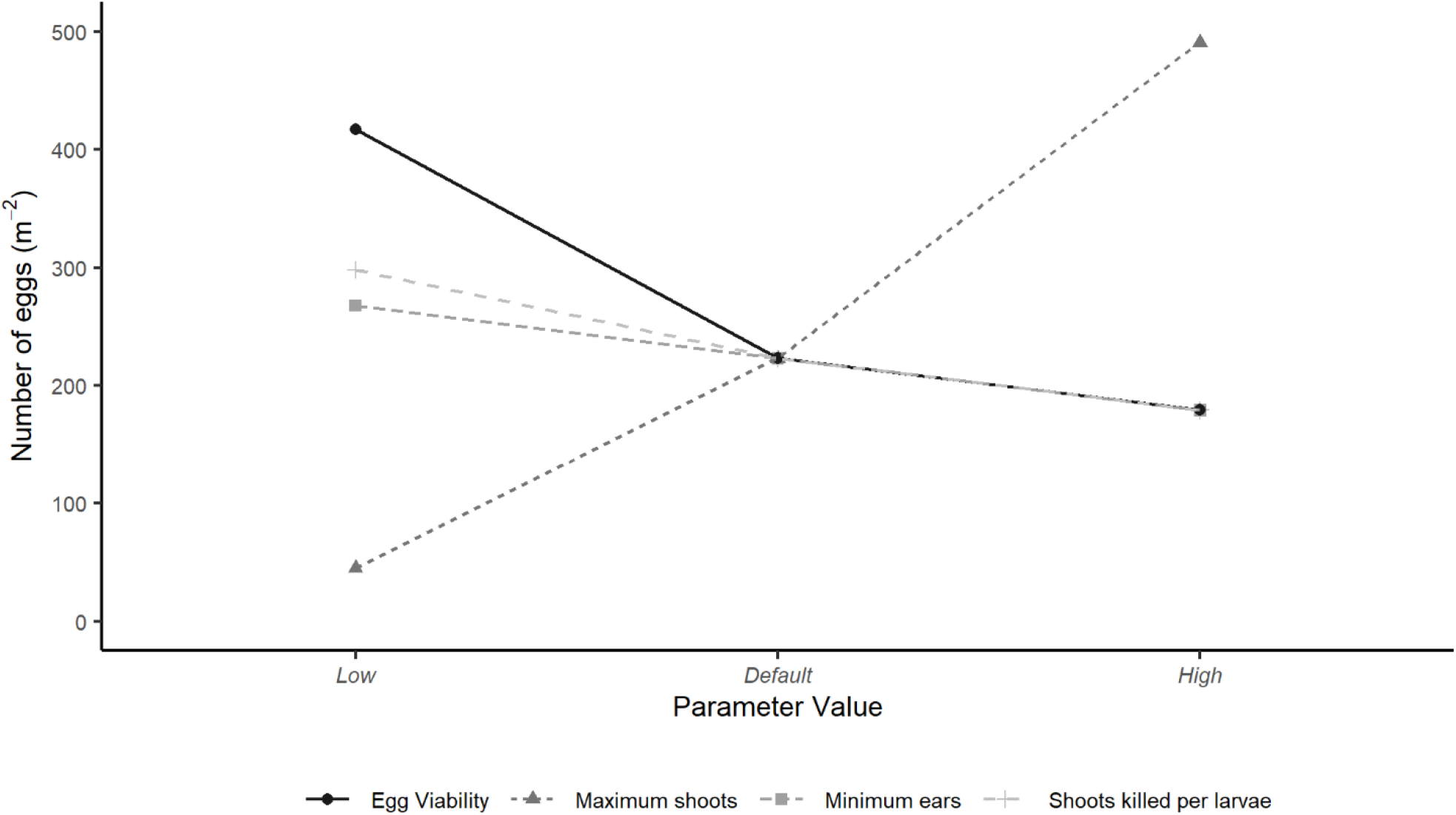
Sensitivity analysis for the economic injury level (EIL) of D. coarctata eggs m^-2^ for the minimum and maximum ranges of each of the parameters used to calculate the EIL. Ranges for each parameter (Low – Default – High): Egg viability = 30%, 56%, 70%; maximum shoots = 600, 1000, 1600, minimum ears = 400, 500, 600; shoots killed per larvae = 3, 4, 5.

The two models we have developed, pest level prediction and crop tolerance prediction (via shoot number estimation), have the potential to represent central components of an IPM strategy for *D. coarctata* control. Although the models have been tested and validated as part of this research, further optimisation is required. For the pest level prediction model this could involve the inclusion of model moderators (soil type, previous crop) to adjust the value using bespoke farm-specific traits. For the shoot number prediction model, this could include the introduction of crop variety and site factors as variables to estimate shoot number using more detailed agronomic factors. The revised threshold scheme will also require experimental validation.

### *A new* D. coarctata *IPM strategy*

In Fig. 9 we outline a *D. coarctata* IPM scenario based on the pest level prediction model, the revised threshold level, and the shoot number prediction model. We believe that this strategy would facilitate non-chemical risk-based control of *D. coarctata* and would comprise the following steps:

1. A seasonal estimation of *D. coarctata* risk per region to advise on the potential level of control required and to enable targeted soil sampling in high-risk regions
2. Use of the revised thresholds to compare predicted *D. coarctata* risk with the minimum number of shoots required to tolerate pest damage while obtaining a viable crop yield
3. Utilisation of the shoot number prediction model to estimate the number of shoots that will be produced for the planned sowing date and sowing rate.
4. Subtraction of the estimated number of shoots required to achieve *D. coarctata* tolerance at the predicted level of *D. coarctata* risk from the estimated number of shoots expected with the planned agronomic practice: A positive value indicates that *D. coarctata* damage can be tolerated naturally, a negative value indicates that additional crop protection or risk-mitigation steps are required, e.g. earlier drilling, higher seed rate, seed treatment (if sown late).
5. The shoot number prediction model can then be used to estimate the minimum plant population required to achieve the minimum shoot number for a given sowing date, or the latest sowing date that could be achieved for a given target plant population.

**Fig. 9:**
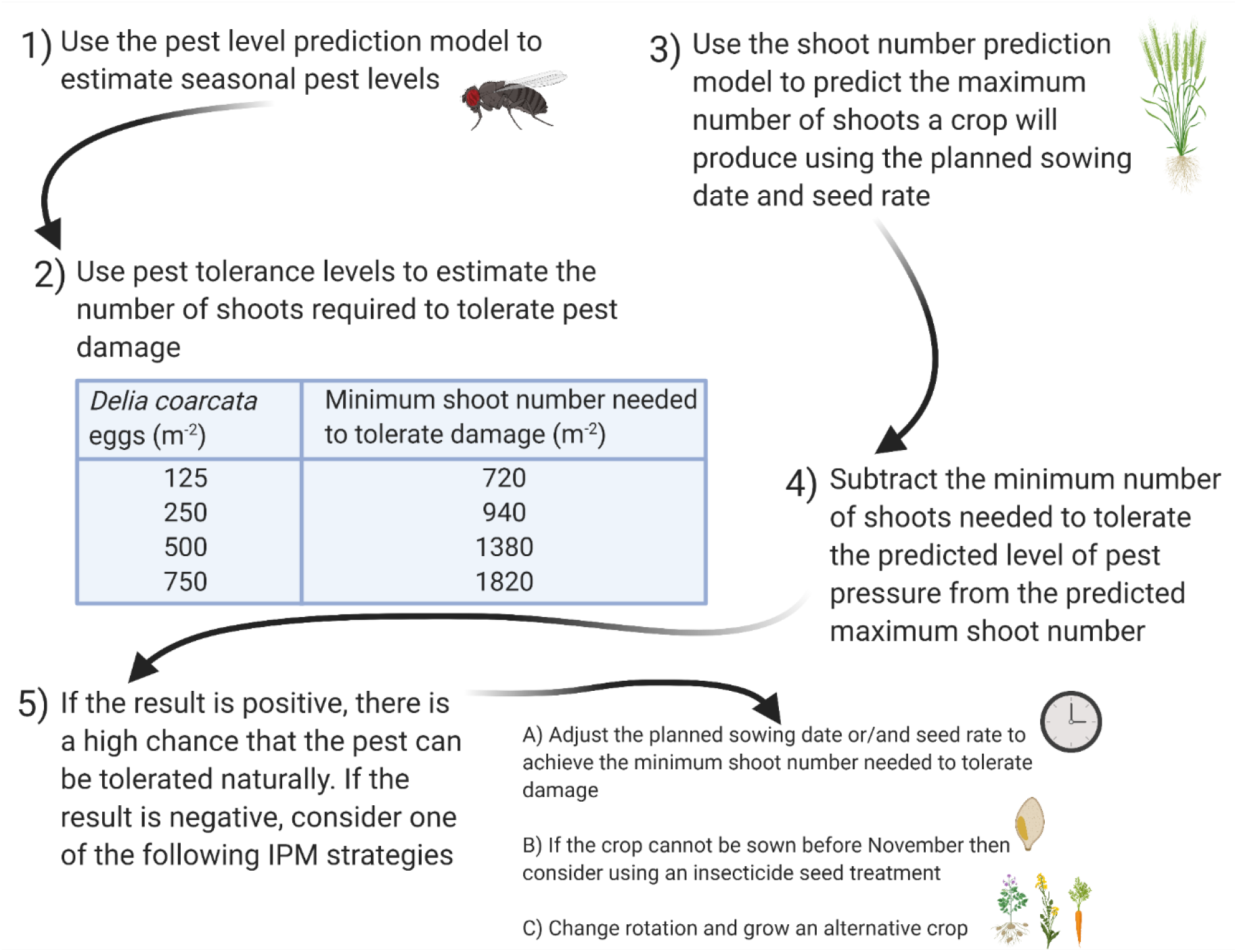
An IPM flow-chart for sustainable management of D. coarctata through the optimisation of pest level and wheat shoot number prediction and crop tolerance – this image was created with BioRender.com

This prescriptive pest management scheme will provide a framework for sustainable *D. coarctata* management. Where the framework indicates that additional crop protection steps are required this could be achieved by adjusting sowing date and/or target plant population to produce a crop with sufficient shoots to tolerate the pest and still achieve potential yield. During seasons of high risk, the option of combining manipulation of sowing date and/or plant population with seed treatment could be employed for late sown crops (November onwards).

## Acknowledgements

The authors gratefully acknowledge the Agriculture and Horticulture Development Board (AHDB) Cereals and Oilseeds for funding this project and would like to thank David Lunn, Josh Humphrey, and Andrew Moore carrying out the 2020 *D. coarctata* surveys and David Lunn, Tom Whiteside, and Nicola Rochford for managing the field trials.

