## Supplementary for "Development of a pest threshold decision support system for minimising damage to winter wheat from wheat bulb fly, *Delia coarctata*"

**Supplementary Material**

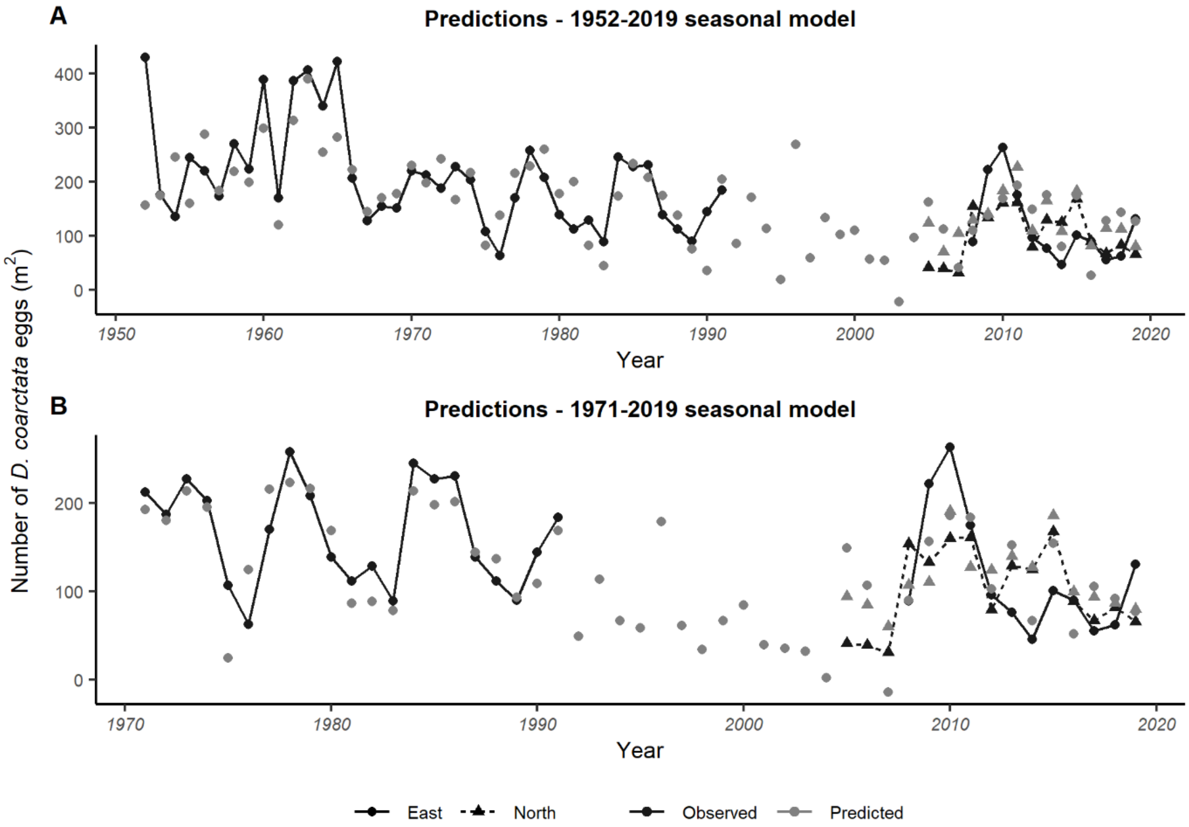


**Fig. S1**: Predicted and observed *D. coarctata* egg numbers. A) 1952 – 2019 seasonal model, B) 1971 – 2019 seasonal model. Predictions (grey) are plotted alongside the mean observed value (black) and divided into the two regions, north (triangle; dashed line) and east (circle; solid line). For clarity, trendlines are included for the observed values only.

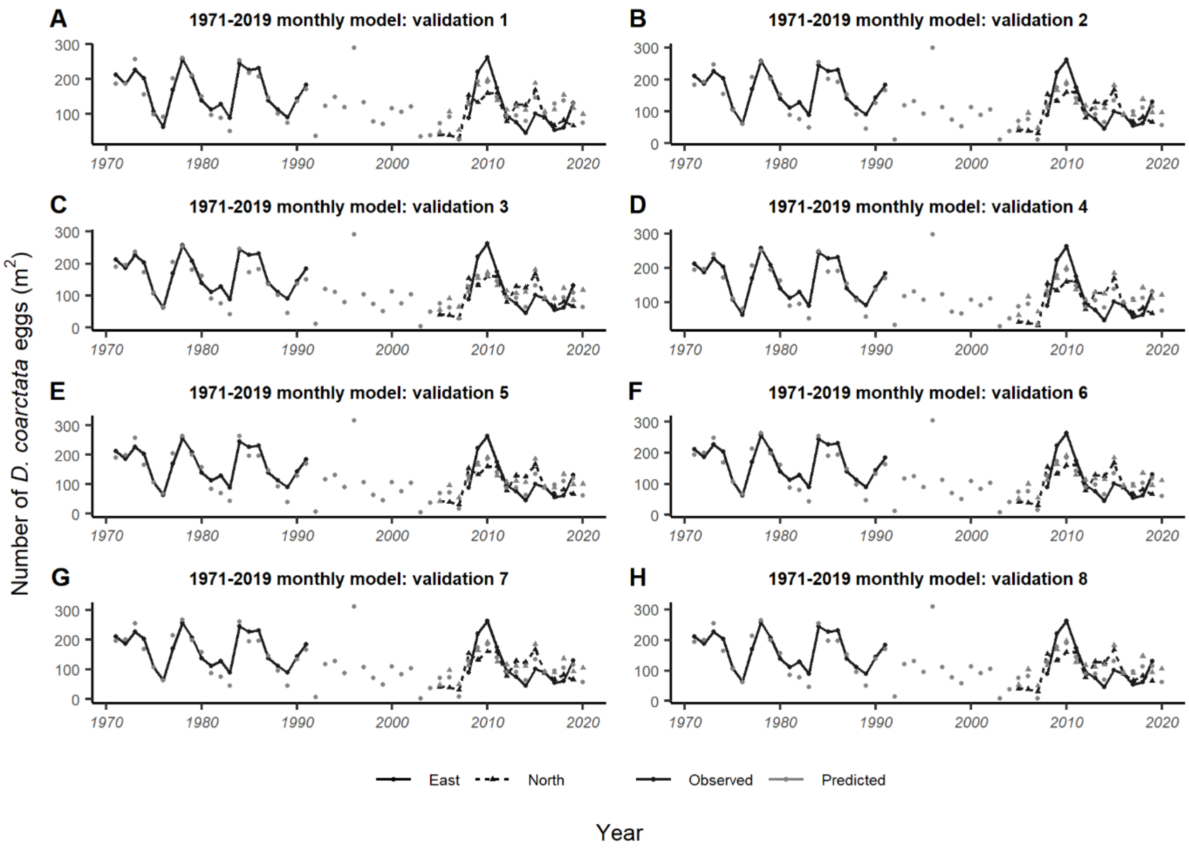


**Fig. S2:** Validation of the 1971 – 2019 monthly model. Graphs show the predicted and observed *D. coarctata* egg numbers for 1971 – 2019 for north (triangle; dashed line) and east (circle; solid line) regions. A) Validation 1 excluding years 1971 – 1975. B) Validation 2 excluding years 2015 – 2019. C) Validation 3 excluding years 1978, 1984 - 1986, 2010. D) Validation 4 excluding years, 2005 - 2007, 2014, 2017. Validations E-H display the validations with five random years removed. For clarity, trendlines are included for the observed values only.

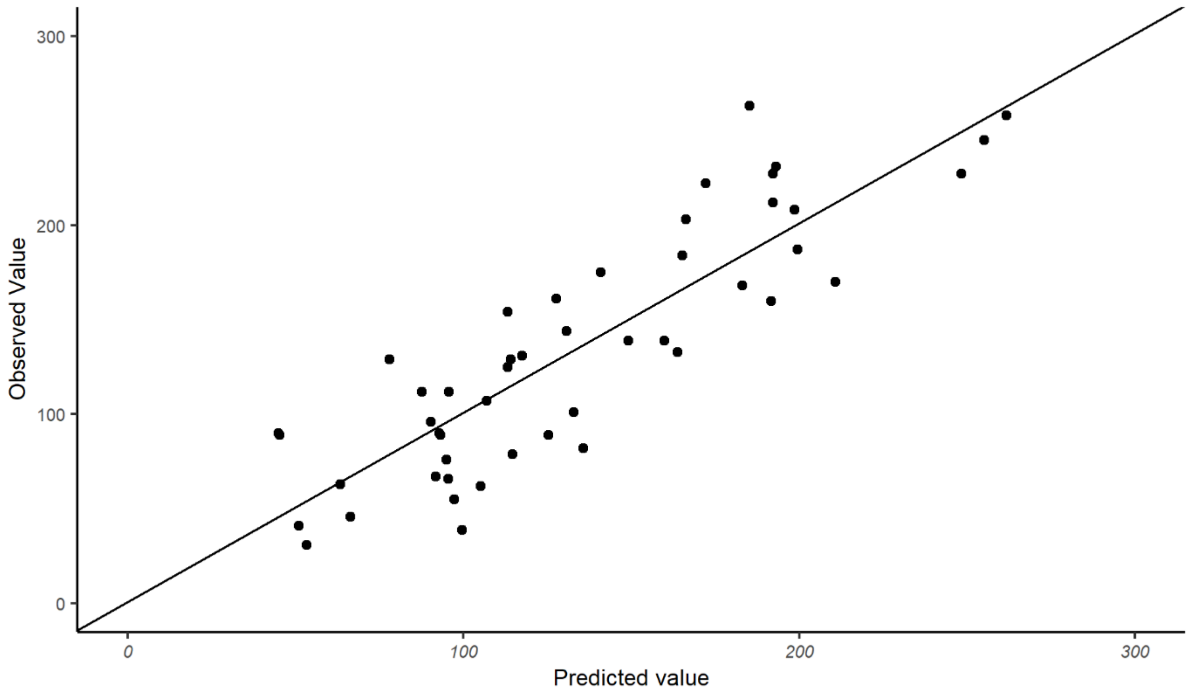


**Fig, S3:** Relationship between the observed vs predicted values for the final *D. coarctata* population prediction model (1971 – 2019 monthly model). Solid line represents a 1:1 line.

**Table S1:** Data on the observed shoot number per plant from the five field trials used to calibrate the shoot number model

| Trial location and harvest year | UK OSM Grid Reference | Sow date (regional timing) | Number of replicates | Seed rate (seeds sown per $m^{-2}$) | | | | | | P Value | SED | Residual df |
|---|---|---|---|---|---|---|---|---|---|---|---|---|
| | | | | 40 | 80 | 160 | 320 | 480 | 640 | | | |
| | | | | Mean shoots per plant at GS31 | | | | | | | | |
| Towthorpe, East Yorkshire, 2016 | SE 90944 62406 | 02/10/15 (standard) | 4 | 6.7 | 5.0 | 7.0 | 4.7 | 3.6 | 3.4 | <0.01 | 0.82 | 15 |
| Huggate, East Yorkshire, 2016 | SE 91669 56414 | 10/10/15 (standard) | 3 | 24.6 | 17.4 | 10.2 | 5.4 | 4.1 | 3.4 | <0.001 | 2.64 | 10 |
| Rosemaund, Herefordshire, 2016 | SO 55793 48437 | 20/09/15 (standard) | 4 | 5.4 | 4.8 | 4.7 | 3.9 | 5.7 | 4.4 | NS | 0.96 | 15 |
| Foxholes, North Yorkshire, 2017 | TA 01400 72200 | 24/10/16 (standard) | 3 | 7.2 | 7 | 5.9 | 4.3 | 3.3 | 3.1 | <0.01 | 0.85 | 10 |
| Bardwell, Suffolk, 2017 | TL 93095 73956 | 29/09/16 (standard) | 3 | 7.7 | 8.6 | 5.4 | 4.3 | 3.7 | 2.5 | <0.001 | 0.75 | 10 |

*All experiments were treated with chlorpyrifos insecticide under an experimental approval.

**Table S2:** Data on the observed shoots per metre squared from the three field trials used to test the shoot number model

| Trial location and harvest year | UK OSM Grid Reference | Sow date (regional timing) | Number of replicates | Insecticide treated* | Seed rate (seeds sown per $m^{-2}$) | | | | | | P Value | SED | Residual df |
|---|---|---|---|---|---|---|---|---|---|---|---|---|---|
| | | | | | 40 | 80 | 160 | 320 | 480 | 640 | | | |
| | | | | | Mean shoots per $m^2$ at GS31 | | | | | | | | |
| Huggate, East Yorkshire, 2016 | SE 91669 56414 | 10/10/15 (standard) | 3 | Untreated | 849 | 919 | 996 | 897 | 845 | 1057 | NS | 105.5 | 10 |
| Foxholes, North Yorkshire, 2017 | TA 01400 72200 | 16/11/16 (late) | 3 | Treated | 12 | 42 | 73 | 159 | 297 | 260 | <0.01 | 54.6 | 10 |
| Bardwell, Suffolk, 2017 | TL 93095 73956 | 25/10/16 (late) | 3 | Treated | 40 | 69 | 76 | 157 | 210 | 262 | <0.01 | 50.2 | 8 |

*Insecticide treated plots were treated with chlorpyrifos insecticide under an experimental approval.
